# Fractional Anisotropy as a Surrogate Marker of Brain Mechanics

**DOI:** 10.64898/2026.04.13.717439

**Authors:** Stefan Rampp, Silvia Budday, Nina Reiter, Nicole Tueni, Jan Hinrichsen, Lars Bräuer, Friedrich Paulsen, Oliver Schnell, Guillaume Flé, Frederik Bernd Laun, Arnd Doerfler

**Affiliations:** Department of Neuroradiology University Hospital Erlangen, Germany; Department of Neurosurgery, University Hospital Erlangen, Erlangen, Germany; Department of Neurosurgery, University Hospital Halle (Saale), Halle (Saale), Germany; Institute of Continuum Mechanics and Biomechanics, Friedrich-Alexander-Universität Erlangen-Nürnberg (FAU), Fürth, Germany; Institute of Functional and Clinical Anatomy, Friedrich-Alexander-Universität Erlangen-Nürnberg (FAU), Erlangen, Germany; Institute of Radiology, University Hospital Erlangen, Friedrich-Alexander-Universität Erlangen-Nürnberg (FAU), Erlangen, Germany

**Keywords:** diffusion weighted imaging, fractional anisotropy, magnetic resonance imaging, mechanics, stiffness

## Abstract

Understanding the mechanical properties of brain tissue may provide crucial insights into brain development, injury, disease and surgical planning. Conventionally, these properties are measured *ex vivo* or in vivo during surgical procedures, while non-invasive *in vivo* alternatives are sparse. This study investigates whether fractional anisotropy (FA) derived from diffusion-weighted magnetic resonance imaging can serve as a surrogate marker for brain tissue stiffness in healthy human brains.

MRI data were collected from three body donor brains, 28 healthy adults, and *a publicly available independent dataset of 26 adults*. FA values were compared with mechanical properties from *ex vivo* mechanical testing of brain tissue.

Statistical analysis revealed a strong negative correlation between FA and the mechanical response for small strains expressed as shear modulus of a one-term hyperelastic Ogden model, indicating that higher FA values are associated with lower tissue stiffness. The nonlinearity parameter alpha exhibited a qualitatively similar, but considerably weaker correlation with FA. These findings were consistent across datasets.

The findings suggest that FA can be a robust, non-invasive marker for estimating mechanical properties of brain tissue, with potential applications in clinical diagnosis and computational modeling of brain mechanics and the study of brain development. Further research is needed to clarify the relationship in lesional tissues and to optimize clinical utility.

## Introduction

Mechanical properties have been shown to play a pivotal role in animal and human brain development, for example the formation of the characteristic cortical patterns of gyri and sulci and interference with these processes may lead to cortical malformations (Budday et al., 2015). Other pathological conditions further demonstrate the significance of tissue mechanics. In traumatic brain injury (TBI), the elastic response of brain tissue to a physical impact defines the location and extent of brain damage, resulting in the typical contre-coup lesions opposite to the initial impact (Orr et al., 2024). Alterations of brain stiffness occur with increasing age, but especially in neurodegenerative diseases (Antonovaite et al., 2021; Sack et al., 2009). Mechanical properties of tumors and their immediate environment can influence the surgical complexity to remove them. Finally, realistic modeling of mechanical brain behavior is necessary to compensate brain shift for neuronavigation and to implement surgical training models (Delorme et al., 2012; Luo et al., 2017).

The (de-facto) gold standard to measure such mechanical tissue properties is testing with specialized equipment, e.g. rheometers and indenters. Depending on the device and the application, the tissue is compressed, sheared or extended and the mechanical response is measured. Obviously, such procedures can only be conducted *ex vivo*, i.e. testing resected tissue or tissue from body donors. *In vivo* application, however, is generally not viable, due to the inaccessibility of sufficient brain tissue and the unavoidable trauma that these devices would induce.

An *in vivo*, non-invasive method to estimate mechanical properties is elastography. The technique induces subtle mechanical deformations, e.g. with vibrations and simultaneously records the minute tissue movements. From these movements, local mechanical tissue properties, i.e. measures of stiffness, can be estimated. One of the first clinical applications was the investigation of liver tissue, e.g. in patients with liver cirrhosis using vibration transducers and ultrasound. In recent years, the technique has been adapted to MRI. Here, extent and direction of subtle tissue movement also induced by vibration is acquired by so called motion-encoding gradients (MEG) with sequences that are exactly synchronized to the frequency of the vibrations. Different inverse methods then allow estimation of tissue properties within each voxel of the scanned volume. With MR-elastography (MRE), investigation of the brain became viable, as ultrasound cannot penetrate the enclosing skull sufficiently.

While MRE is a comparably novel, non-invasive method to probe mechanical properties of the brain, there are currently some limitations for clinical application. For instance, the choice of parameters, e.g. vibration frequencies and duration but also MR sequence characteristics, is known to influence the stiffness estimate. Furthermore, deriving local mechanical properties from tissue movement requires the solution of an inverse problem that depends on underlying modeling assumptions, thereby introducing additional degrees of freedom. On the other hand, *ex vivo* validation data are scarce and largely restricted to samples obtained from surgical resections or body donors, which are also used for that direct mechanical testing.

A further non-invasive avenue to *in vivo* estimation of mechanical properties may not be their direct measurement but the investigation of their structure correlates, i.e., the underlying histological architecture which may give rise to tissue stiffness. Previous studies indicate that MR diffusion-weighted imaging (DWI) may provide such surrogates. For example, some studies demonstrated that fractional anisotropy (FA) correlates with the stiffness of meningiomas experienced intraoperatively (Bao et al., 2024; Kashimura et al., 2007).

FA reflects the local structure of diffusion, i.e., mainly the local architecture of fiber tracts. FA is low in locations with a predominant fiber orientation, e.g., in the corpus callosum or the internal capsule, while spaces with crossing fibers would result in high FA values. Hypothetically, anisotropy, i.e. locally concentrated heterogeneous fiber directions would correspond to mechanical stiffness, analogous to timber framework in building construction, where angled, crossing beams provide static stability.

In the current study, we evaluate whether MR-FA correlates with mechanical stiffness in normal brain tissue, i.e., under physiological conditions, in contrast to previous work available in the literature. We compare parameters of a hyperelastic mechanical model determined from body donor brain tissue via *ex vivo* testing with MR-FA-data from body donors and healthy controls. The results could be used to further optimize and validate MRE, refine computational models and support clinical diagnosis of, e.g., cortical malformations, tumors and neurodegenerative diseases.

## Methods

### Participants

#### Healthy participants

A total of 28 healthy participants were included in the study. Inclusion criteria were adult age, MR compatibility and absence of any brain pathologies or history thereof. Mean age was 30 ±9.7 years (range 19 to 57 years); 17 (61%) were women. Data was acquired during two separate studies, which were reviewed and approved by the local institutional review board (registration numbers 23-389-Bm and 24-477-Bm) and which included the agreement to utilize the data for related research. Written informed consent was obtained from all participants.

#### Independent dataset

To investigate reproducibility of the findings and demonstrate independence from the specific scanner model, analyses were additionally performed using a public dataset (Tian et al., 2022). The study included 26 cognitively normal adults with a mean age of 36.8 ± 14.6 years (range 22-72 years); 17 participants were women (65%).

#### Body donors

For intraindividual regional comparison of FA and mechanical properties from *ex vivo* testing, three body donor brains were included. Body donor 1 was a 71-year-old male. The cause of death was acute liver and heart failure. Post-mortem time was 55h. Pathologies of the brain were not reported. Body donor 2 was a 57-year-old male. Cause of death was renal failure caused by a tumor lysis syndrome due to a neuro-endocrine tumor. The exact time of death is unknown, post-mortem time was more than 48 hours. Hypoxic/ischemic infarction of the right basal ganglia and cerebral micro/macroangiopathy were reported. Body donor 3 was a 60-year-old female who died of multi-organ failure after hemorrhagic shock and acute respiratory distress syndrome due to pneumogenic sepsis. Post-mortem time was 39 hours. Brain pathologies were not reported.

Procedures were reviewed and approved by the local institutional review board (registration numbers 22-209-Bp).

### MRI acquisition

Due to logistical constraints and different acquisition time points before (body donors) and after a system upgrade (controls), MRI data was acquired with different scanners.

#### Healthy participants – inhouse dataset

Diffusion images were acquired on a 3T scanner (Magnetom Cima.X, Siemens Healthineers, Forchheim, Germany) using a 2D EPI diffusion sequence with TE=44 ms, and TR=3500 ms. The slice thickness was 2 mm. The spacing between slices was 2.6 mm. The in-plane resolution was 1.71875 mm x 1.71875 mm. The flip angle was 90 degrees. The pixel bandwidth was 1698 Hz. A DTI diffusion scheme was used, and a total of 30 diffusion sampling directions were acquired. The b-value was 1000 s/mm_2_. The tensor metrics were calculated using DWI with b-value lower than 1750 s/mm_2_.

In addition, an 3D-T1-weighted MPRAGE was acquired with an isotropic voxel resolution of 1 mm (TI = 900 msec, flip angle 9°).

#### Healthy participants – independent dataset

For details on acquisition and data processing of the independent dataset, see (Tian et al., 2022). In short, data were acquired on a 3 T Connectome scanner (Magnetom CONNECTOM, Siemens Healthineers, Erlangen, Germany) using a 2D EPI diffusion sequence with TE=77 ms and TR = 3800 ms. The in-plane resolution was 2 mm x 2 mm, slice thickeness was 2 mm. A total of 16 different B-values (50-6000 s/mm^2^) were used with 16 different encoding directions for b < 2400 s/mm2 and 64 directions for b ≥ 2400 s/mm^2^. Data was subjected to motion and image distortion correction.

A 3D-T1-weighted MPRAGE with an isotropic voxel resolution of 1 mm (TI = 1100 ms, flip angle 7°) was acquired in addition.

DTI reconstruction and FA calculation was conducted using DSI Studio (Yeh, 2025) (accessed May 2024, http://dsi-studio.labsolver.org).

#### Body donors

Body donor brains were kept unfixated to allow for mechanical *ex vivo* testing after the MRI scans. To slow post-mortem tissue decomposition, brains were kept in a transport cooler at approximately 8-10°C.

The brains were scanned embedded in artificial CSF (see suppl. material) within 3D-printed container devices without additional cooling. For scans of brains 1 and 3 a device according to (Kim et al., 2021), and for brain 2 a customized container was used.

The diffusion images of body donor brain 3 were acquired with the same approach as the inhouse healthy controls. Body donor brains 1 and 2 were acquired on a 3T Magnetom Prisma scanner (Siemens Healthineers, Erlangen, Germany) using a 2D EPI diffusion sequence with TE=69 ms, and TR=4100 ms. The slice thickness was 4 mm and the spacing between slices was 5.2 mm. The in-plane resolution was 1.71875 mm x 1.71875 mm. The flip angle was 90 degrees. The pixel bandwidth was 1955 Hz. A DTI scheme was used, and a total of 20 diffusion sampling directions were acquired at a b-value of 1000 s/mm_2_.

### MRI processing

Segmentation of the T1w images was performed with the Computational Anatomy Toolbox (CAT12) (Gaser et al., 2024) included in Brainstorm (Tadel et al., 2011), which is documented and freely available for download online under the GNU general public license (http://neuroimage.usc.edu/brainstorm). The segmentation procedure fits several anatomical atlases and parcellation schemes to the individual MRI. From these, the Hammers (Hammers et al., 2003), Mori (Oishi et al., 2009) and IBSR atlas (*Internet Brain Segmentation Repository*, 2013), as well as the gray and white matter compartments from the CAT12 tissue classifications were used to intraindividually define the governing regions from Hinrichsen et al. (Hinrichsen et al., 2023) (table 1).

**Table 1:** Governing regions and abbreviations. Region MT summarizes midbrain and thalamus due to similar mechanical behavior, referred to with the abbreviation ‘M’ in (Hinrichsen et al., 2023).

| Governing region | Abbreviation |
| --- | --- |
| Corpus callosum | CC |
| Corona radiata | CR |
| Cerebellum | CB |
| Hippocampus | Hi |
| Midbrain and thalamus | MT |
| Brainstem | BS |
| Basal ganglia | BG |
| Amygdala | Am |
| Cortex | C |

**Table 2:**
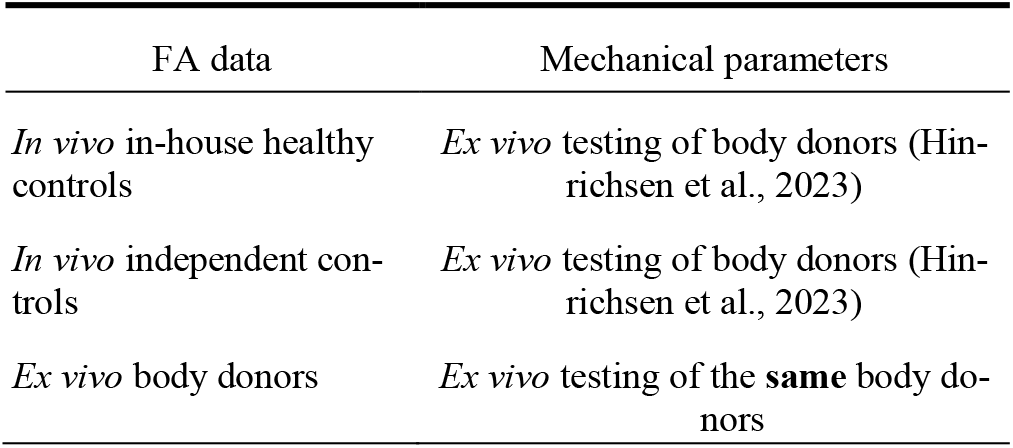
Overview of included datasets and comparison values from mechanical *ex vivo* testing.

| FA data | Mechanical parameters |
| --- | --- |
| *In vivo* in-house healthy controls | *Ex vivo* testing of body donors (Hinrichsen et al., 2023) |
| *In vivo* independent controls | *Ex vivo* testing of body donors (Hinrichsen et al., 2023) |
| *Ex vivo* body donors | *Ex vivo* testing of the **same** body donors |

In body donor brains, automated segmentations were not always acceptable e.g., due to deformations and susceptibility artifacts. In these cases, regions were manually segmented, some partially and excluding areas affected by artifacts.

Subsequently, the average FA value was calculated over all voxels within each region, including left and right homologues of regions present bilaterally.

### Mechanical testing

Mechanical properties of body donor brain tissue were measured using the procedure described in Hinrichsen et al. (Hinrichsen et al., 2023). In short, using a biopsy punch, cylindrical samples of 8 mm diameter and approximately 5 mm thickness were extracted from different regions. Tissue response under compression, tension and torsional shear was measured with a Discovery HR-3 rheometer (TA instruments, New Castle, Delaware, USA). A hyperelastic one-term Ogden model was then fitted to the preprocessed and averaged data (neglecting poro- and viscoelastic effects) using an iterative inverse parameter identification procedure. In samples with different loading directions during testing, e.g., perpendicular and parallel to the main fiber direction in the corpus callosum, resulting parameters were averaged.

For evaluation of data in healthy participants, *ex vivo* mechanical properties of each governing region were taken from Hinrichsen et al. (Hinrichsen et al., 2023), which were determined by applying the above procedure to 182 specimens from seven human brains.

Mechanical parameters were available for the unconditioned and preconditioned responses, represented by the first and third loading cycles, respectively, as well as for two distinct Poisson’s ratios. Poisson’s ratio is a measure of the transverse deformation occurring perpendicular to the direction of applied loading. It characterizes to the material’s compressibility, as higher values approaching 0.5 indicate near-incompressibility, while lower values reflect greater volume changes. It was set to ν = {0.49, 0.45}. Trends persisted over all four datasets, and we chose to display data for the unconditioned response and for ν = 0.45.

### Statistical evaluation

Statistical evaluation of FA vs. mechanical properties (shear modulus and nonlinearity parameter alpha) was conducted separately for the three groups: In-house healthy participants, independent dataset of healthy participants and body donors. Correlation between the shear modulus, respectively alpha and FA values was evaluated by repeated measures correlation, controlling for subject effects. To evaluate consistency of regional FA, shear modulus and alpha values between datasets, (Pearson) correlation between regional means over all subjects of the group was calculated. For comparison of FA variability between in-house and independent dataset, regional interquartile FA ranges were subjected to a two-sided paired t-test. All statistical analyses were conducted with R version 4.5.2 (R Core Team, 2025) and the rmcorr (version 0.7.0) package (Bakdash & Marusich, 2017).

## Results

### Inhouse healthy controls

FA in inhouse healthy controls, calculated with manufacturer’s software, showed a strong correlation with the shear modulus derived from *ex vivo* mechanical testing of body donor samples (figure 1). The repeated measures correlation analysis yielded r = - 0.808 (95% confidence interval -0.849 - -0.757, p < 0.0001, fig. 1a), i.e., higher FA values were associated with lower shear modulus values. Mean FA in the cerebellum and hippocampus deviated slightly to lower FA values, whereas the brainstem showed weakly elevated FA.

**Figure 1:**
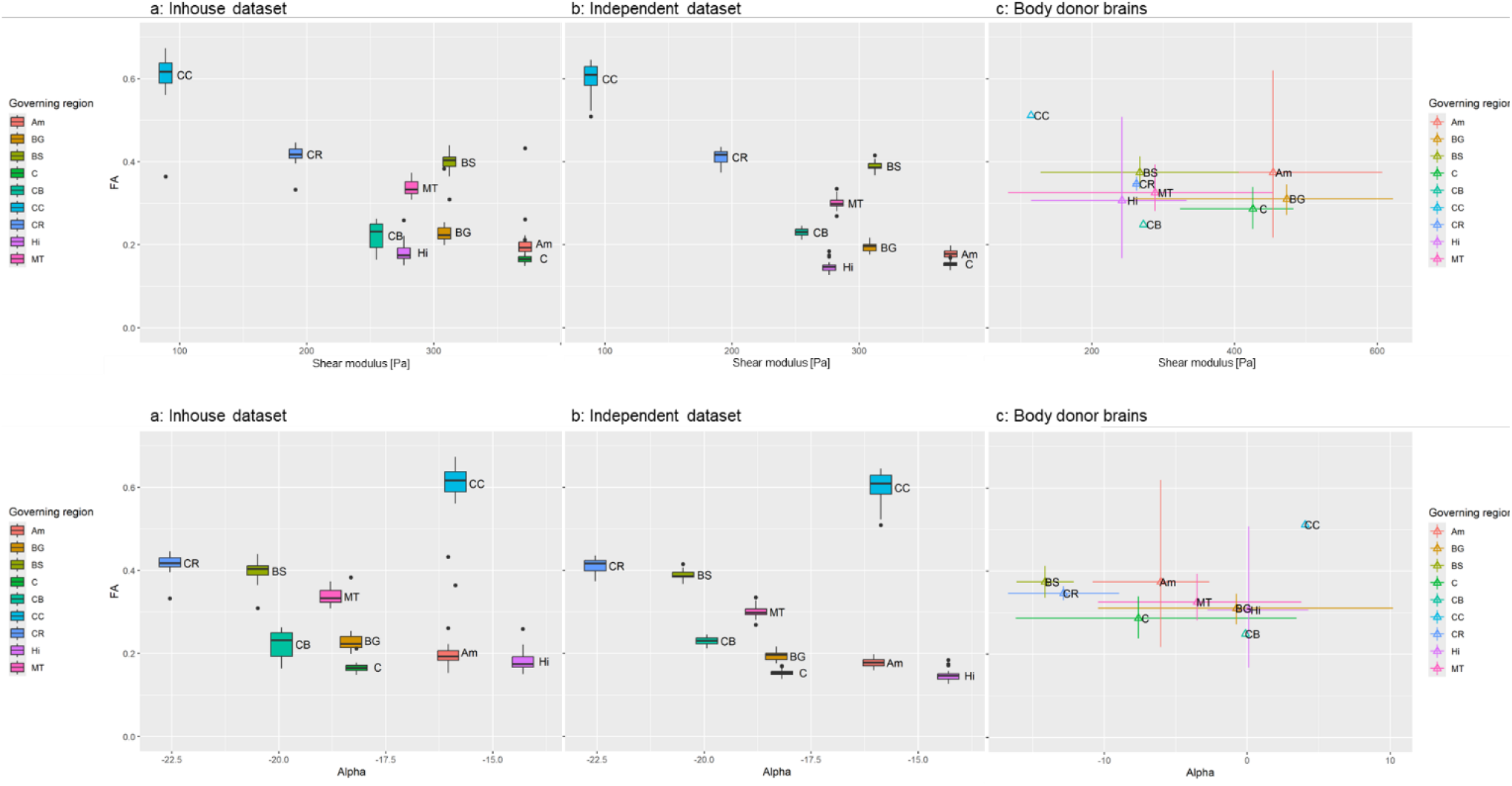
FA-values compared to shear modulus (top row) and alpha (bottom row) in the three datasets: a: in-house *in vivo* healthy controls, b: independent dataset of *in vivo* healthy controls, and c: three body donor brains. Mechanical properties were taken from (Hinrichsen et al., 2023) for datasets a and b, and from mechanical testing of the body donor brains for dataset c. Box-and-whiskers plots show medians, first and third quartile. The whiskers extend from the first/third quartile to the largest value no further than 1.5 * interquartile range from the first/third quartile. More extreme values are depicted as dots. Plot c shows mean values and ranges for each available region.

In contrast, FA showed only a weak but statistically significant correlation to the nonlinearity parameter alpha (r = -0.183, -0.306 - -0.053, p = 0.006). Higher FA values were related to low/more negative alpha values. Here, the most extreme outlier was the corpus callosum, which showed the highest observed FA values for high alpha values.

### Independent dataset of healthy controls

Results of the independent dataset were highly compatible with the inhouse data. Repeated measures correlation between FA and shear modulus was r = - 0.830 (-0.868 - -0.782, p < 0.0001). In this dataset also, the cerebellum and hippocampus exhibited comparatively lower FA values, while the brainstem showed higher FA values, consistent with expectations based on the shear modulus.

FA and alpha also showed a considerably weaker correlation (r = -0.232, -0.356 - -0.099, p = 0.0007) and the corpus callosum as outlier.

Notably, FA values for the individual regions were highly similar to the inhouse dataset.

### Body donor brains

The mechanical properties of body donors, as measured by shear modulus and nonlinearity alpha, showed an overall shift toward higher values compared to (Hinrichsen et al., 2023). Alpha even reached positive values in the corpus callosum, indicating a pronounced tension response. The corona radiata and basal ganglia showed higher shear modulus values in the interregional comparison, while the opposite trend was observed for the cortex and hippocampus. In the interregional comparison, the hippocampus, amygdala, brain stem, and cortex ranked lower.

Still, the inverse relationship of FA and shear modulus could be reproduced in body donors in principle, however with limitations. Overall, both shear moduli and FA values showed considerable variation. Inspection of the FA data revealed distortion artifacts due to remaining air bubbles and vicinity to the container wall especially in basal areas, i.e., the hippocampus and amygdala, which showed the largest variability. Including all areas in the statistical evaluation did not yield a significant correlation of FA and shear modulus (r = -0.116, -0.542 – 0.357, p = 0.63) or alpha (r = -0.142, -0.560 – 0.334, p = 0.56). Exclusion of hippocampus and amygdala however results in a significant correlation of FA and shear modulus (r = -0.584, -0.859 - - 0.049, p = 0.03), but not for FA and alpha (r = 0.018, -0.537 – 0.563, p = 0.95).

### Consistency between datasets

Ranges of FA, shear modulus and alpha values were overall consistent with the control datasets in most areas, correlations between groups ranged between 0.755 and 0.995 for the different parameters with all correlations showing statistical significance (p < 0.05). Especially the more extreme results of the corpus callosum with high FA, low shear modulus but high alpha could be reproduced. FA values of the independent dataset showed significantly lower variability than the inhouse data (p = 0.026, two-sided paired t-test of regional interquartile FA ranges).

## Discussion

Evaluation of FA derived from diffusion tensor imaging demonstrated a strong negative correlation with the shear modulus determined by *ex vivo* mechanical testing of brain tissue: Higher FA values indicated a lower regional stiffness. A correlation with the non-linearity parameter alpha was considerably weaker but detectable and showed the same direction. FA reflects the local microstructural fiber architecture. High values are encountered in cable-like structures, e.g. the pyramidal tract or the corpus callosum. Low FA values are associated with differently oriented fibers within the observed voxel without a dominant direction, e.g. in nuclei and gray matter.

The distribution of forces across reinforced structural elements with varying orientations is a fundamental principle in architecture, and has long been employed, e.g., in traditional timber framing. The different beams and frames transfer forces from different directions and redistribute them to avoid mechanical deformation or damage. The same architectural principle may also have relevance in the context of fiber microstructure of the brain.

### FA and mechanical properties

Previous studies have explored the correlation of FA with mechanical stiffness. Kashimura et al. (Kashimura et al., 2007) utilized FA to differentiate subtypes and intraoperatively determined consistency of meningiomas to support planning of surgery. They observed higher FA in fibroblastic vs. meningothelial, as well as in hard vs. soft tumors. While their findings support the association of FA with measures of stiffness, our results show the opposite direction, i.e., high FA was related to low stiffness as expressed by low shear modulus values. Two main factors may explain these seemingly contradictory results. First, Kashimura et al. determined stiffness with intraoperative evaluation by the surgeon, which may not directly correspond to *ex vivo* rheometer testing. The latter measures the tissue sample in isolation, i.e. tissue environment, CSF, blood pressure and other relevant factors *in vivo* (Herthum et al., 2021) are not considered. Second, our study focused on the evaluation of healthy tissue. It seems conceivable that mechanical behavior of lesional tissue is impacted by further factors, e.g. pathological growth, scarification and calcification.

Bao et al. (Bao et al., 2024) showed that among several techniques, the combination of MR elastography (MRE) and FA provides the best differentiation between hard and soft meningiomas (area-under-the-curve (AUC) value 0.88). MRE alone achieved a slightly lower performance (AUC 0.81), while FA alone showed more limited but still significant differentiation (AUC 0.66). Interestingly, in their study also, FA was increased in harder tumors. The authors further confirmed these results with *ex vivo* indentation tests, ruling out relevant influences of subjective evaluation or impact of the anatomical environment. Still, an evaluation of healthy tissue could not be performed.

In contrast, Cepeda et al. (Cepeda et al., 2021) demonstrated a strong positive correlation of FA with the mean tissue elasticity (MTE) (with behavior inverse to stiffness) determined by intraoperative ultrasound elastography (USE) of low- and high-grade glioma but also meningioma. They also reported significantly different intratumoral FA values with meningioma showing the highest values, followed by low- and then high-grade glioma with concordantly associated MTE data. Overall, their findings correspond well with our comparison with *ex vivo* rheometer testing. The discrepancy to Bao et al. and Kashimura et al. however remains unresolved.

Our results also suggest a correlation with the non-linearity parameter alpha. This material parameter of the one-term hyperelastic Ogden model represents how the tissue stiffens or softens under deformation. Negative values as mostly applied here indicate stiffening with increasing compression. Nevertheless, differences may influence the subjective impression of a tissue’s stiffness, e.g., when palpated intraoperatively. It likely also contributes to the location and extent of lesions after traumatic brain injury (TBI). While frequently explored in material sciences (Hinrichsen et al., 2023) and in the context of MRE (Herthum et al., 2021), there are no studies relating FA to alpha.

Our results suggest a negative correlation of FA with alpha, however much weaker than with the shear modulus. The explained variance of the observed data (r^2^) reaches only ∼2-5% in the inhouse and independent datasets. Correspondingly, FA does not describe the nonlinear behavior well and other factors beyond microstructural fiber architecture as imaged by DTI likely play a role. The results of FA and alpha in the corpus callosum are especially interesting. These deviate considerably from the expected correlation trajectory in the same manner in all three datasets (fig. 2). FA values in the corpus callosum are among the highest but at the same time show the largest, i.e., least negative alpha values. The mechanical behavior therefore deviates from that of other regions, such as the corona radiata, which also consist of highly aligned white matter fibers. Histological analysis shows considerable differences (Reiter et al., 2025), which may explain our findings. In the corpus callosum, glial cells and blood vessels show a highly aligned organization in addition to the axons. In the corona radiata, however, glial cells are grouped between crossing axon bundles. Since diffusion along axons provide the dominant contribution to the DWI signal, differences in glial and vascular architecture may not be adequately reflected by FA and may further result in different mechanical properties.

**Figure 2:**
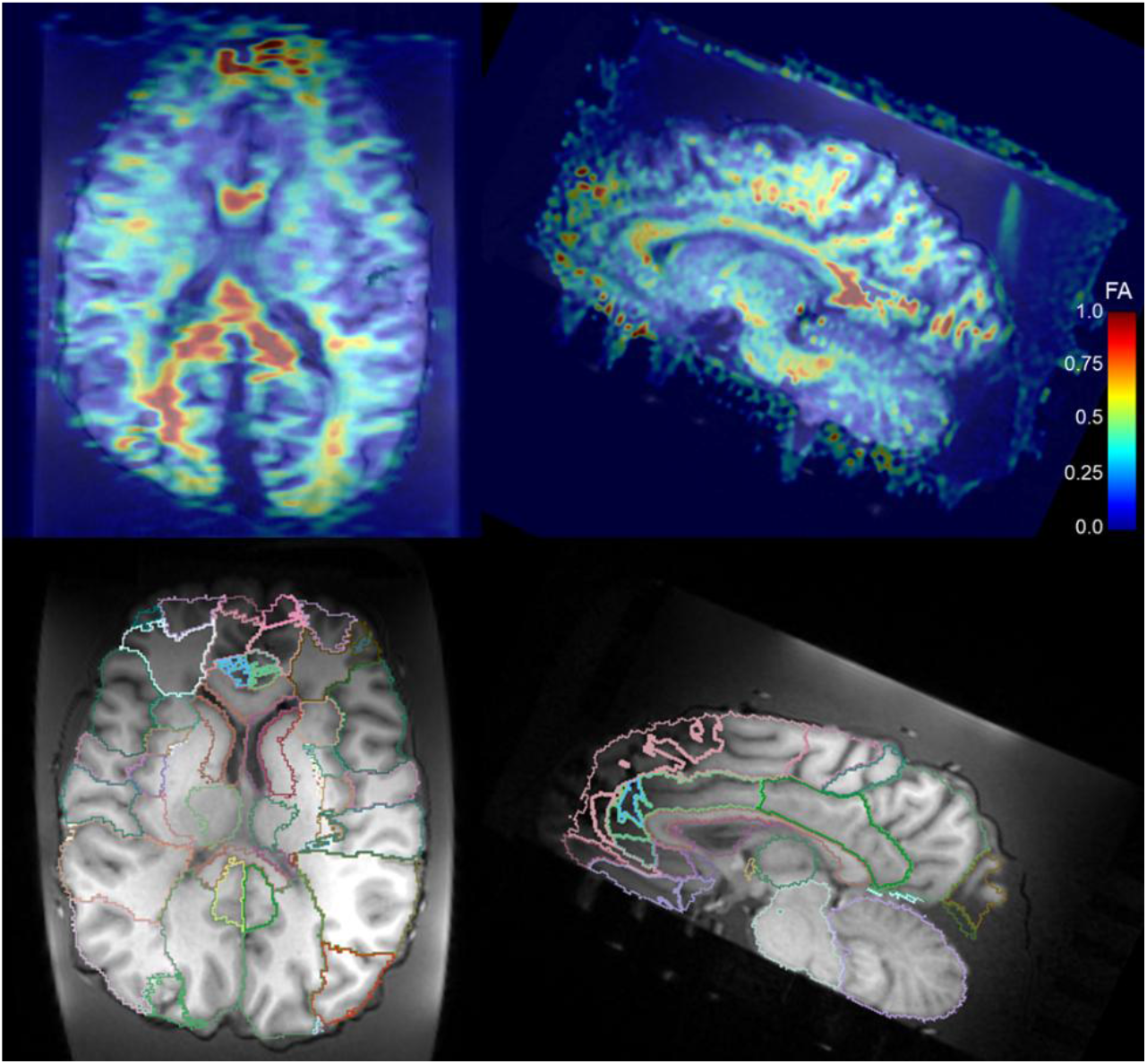
Example of body donor data (body donor 3). Top: FA data overlaid on T1. Bottom: T1 with superimposed Hammers atlas.

### Reproduction in different datasets

The correlation of FA with mechanical properties was reproduced in two independent datasets *in vivo*. Notably, the specific techniques applied for DWI were rather different. While the inhouse series employed the standard manufacturer-provided sequence and reconstructions, the independent data (Tian et al., 2022) utilized a more sophisticated, highly optimized approach. Considerable differences of FA values and their correlations were not observed, underlining the robustness of our principal finding. An effect of optimized DTI however was apparent in the significantly lower regional variability of the FA data. Consequently, exploration of FA for estimation of mechanical properties is likely to benefit from such approaches. An extension to Q-space trajectory imaging (QTI) (Westin et al., 2016) may provide a further level of detail.

The association of FA and shear modulus was also supported by the body donor results. In contrast to the *in vivo* data, regional FA and *ex vivo* mechanical properties could be related intraindividually. The low sample size, potential susceptibility artifacts and acquisition of diffusion below body temperature (Berger et al., 2022) limit the results’ robustness. However, increasing shear modulus values showed the same qualitative decrease of FA as the *in vivo* data. Statistical significance was achieved after excluding two regions affected by susceptibility artifacts.

## Conclusion

Estimating tissue stiffness non-invasively over the whole brain opens new avenues to construct computational models of brain mechanics. FA and MRE, and potentially their combination, could be used to develop personalized models with voxel-specific parameters. Potentially, these could be used to detect pathological changes, e.g. around brain tumors or within subtle epileptogenic lesions, such as focal cortical dysplasia. However, for such clinical applications, a better understanding of how FA and other modalities reflect mechanical properties of lesional tissue is needed.

For the investigation of brain development, FA may be especially interesting due to its applicability already during the prenatal stage (Di Stefano et al., 2025) allowing for intraindividual longitudinal studies.

## Data Availability

The MRI data of the in-house cohort and the body donors are available on Zenodo (https://doi.org/10.5281/zenodo.19454975).

## Acknowledgment

The authors wish to sincerely thank those who donated their bodies to science so that anatomical research could be performed. Results from such research can potentially improve patient care and increase mankind’s overall knowledge. Therefore, these donors and their families deserve our highest gratitude.

We further thank Lisa Stache of the Institute of Functional and Clinical Anatomy for the preparation of the body donor brains.

## Supplementary material

### Composition of artificial CSF used to store body donor brains

#### Ringer’s base solution 1

Ringer's base solution 1
|  |  |  |
| --- | --- | --- |
| NaCl | 124 mM | 72.466 g/l |
| NaH<sub>2</sub>PO<sub>4</sub> | 1.25 mM | 1.725 g/l |
| MgSO<sub>4</sub> | 1.80 mM | 4.437 g/l |
| CaCl<sub>2</sub> | 1.60 mM | 2.352 g/l |
| KCl | 3.0 mM | 2.237 g/l |
| Glucose | 10 mM | 19.817 g/l |

#### Ringer’s base solution 2

Ringer's base solution 2
|  |  |  |
| --- | --- | --- |
| NaHCO<sub>3</sub> | 26 mM | 21.843 g/l |

#### Ringer’s working solution (artificial CSF)

For 1 liter working solution:

100 ml base solution 1 + 100 ml base solution 2 + 800 ml aqua dest.

pH adjusted to 7.4

